# IntelliR: A comprehensive and standardized pipeline for automated profiling of higher cognition in mice

**DOI:** 10.1101/2024.01.25.577156

**Authors:** Vinicius Daguano Gastaldi, Martin Hindermann, Justus BH Wilke, Anja Ronnenberg, Sahab Arinrad, Sabine Kraus, Anne-Fleur Wildenburg, Antonios Ntolkeras, Micah J Provost, Liu Ye, Yasmina Curto, Jonathan-Alexis Cortés-Silva, Umer Javed Butt, Klaus-Armin Nave, Kamilla Woznica Miskowiak, Hannelore Ehrenreich

## Abstract

In the rapidly evolving field of rodent behavior research, observer-independent methods facilitate data collection within a social, stress-reduced, and thus more natural environment. A prevalent system in this research area is the IntelliCage, which empowers experimenters to design individual tasks and higher cognitive challenges for mice, driven by their motivation to access reward. The extensive amount and diversity of data provided by the IntelliCage system explains the growing demand for automated analysis among users. Here, we introduce IntelliR, a standardized pipeline for analyzing raw data generated by the IntelliCage software, as well as novel parameters including the cognition index, which enables comparison of performance across various challenges. With IntelliR, we provide the tools to implement and automatically analyze 3 challenges that we designed, encompassing spatial, episodic-like, and working memory with their respective reversal tests. Using results from 3 independent control cohorts of adult female wildtype mice, we demonstrate their ability to comprehend and learn the tasks, thereby improving their proficiency over time. To validate the sensitivity of our approach for detecting cognitive impairment, we used adult female NexCreERT2xRosa26-eGFP-DTA mice after tamoxifen induced diphtheria toxin-mediated ablation of pyramidal neurons in cortex and hippocampus. We observed deterioration in learning capabilities and cognition index across several tests. IntelliR can be readily integrated into and adapted for individual research, thereby improving time management and reproducibility of data analysis.

**HIGHLIGHTS:**

- IntelliR is a standardized pipeline for analyzing raw data of IntelliCage software.
- Domains include spatial, episodic-like, and working memory with reversals.
- WT mice (3 cohorts) comprehend, learn and improve proficiency over time.
- Cognition index permits comparison of performance across cognitive domains.
- Mice with ablation of pyramidal neurons decline mainly in working memory.

## INTRODUCTION

The rapid development of elaborate mouse models for neuropsychiatric diseases and human brain pathology created a pressing need for comprehensive and well-standardized high-throughput behavioral phenotyping protocols (1, 2). In response, a variety of standardized scoring systems and behavioral testing protocols have been developed, validated, and proven extremely useful. These methods typically rely on either fully automated equipment, semi-automated tracking software, or experimenter-dependent testing and scoring of predefined readouts (2–4). Yet, despite the attempt to standardize these tests, considerable variability and potential stressors may be introduced by environmental factors, singling of mice for testing, handling, and diverse housing conditions (1, 2). To minimize these problems, the IntelliCage system was developed to enable automated phenotyping of mice in an observer-independent social setting, using longitudinal monitoring and cognitive profiling of individual mice in their home cage amidst peers (1, 2, 5) and has shown great reproducibility across different laboratories (6, 7).

An IntelliCage provides space for up to 16 mice and is equipped with 4 computer-controlled corners that function as operant chambers with just enough space for a single mouse. Mice are individually tracked by small subcutaneous radiofrequency identification transponders that are registered by antennas within the corner entries. Each corner is equipped with programmable light-emitting diodes, and two drinking bottles with built-in ‘lickometers’ that register contacts with the drinking nipple. Access to the drinking nipples is provided through holes with nose poke sensors and is controlled by motorized doors (2, 5). The corners can be programmed to restrain or enable access to the bottles based on conditions such as time, location of the corner, presence of an operant response (i.e., nose poke), a sequence of events, and more (1, 2, 8–10). Dependent on the testing protocol, learning can be reinforced either positively via rewards, such as access to tap water, sucrose, drug solutions, or negatively via aversive stimuli, such as airpuffs, bitter, sour or allergen solutions, or by light-emitting diodes that provide diverse visual stimuli (2). This versatility facilitated the development of a multitude of IntelliCage-based challenges assessing diverse behavioral domains, including general and circadian activity, exploration, stereotypy, emotionality, social behavior, consumption behavior (i.e., preference/avoidance of drugs or allergens), and finally learning and memory (2). However, despite the increasing number of available IntelliCage-based challenges, data analyses and statistical methods vary considerably across studies (2, 5, 7, 11–18) and often rely on manual data extraction from the graphical user interface of the IntelliCage Analyzer software, which is time consuming, and error-prone (18).

To improve reproducibility and to overcome the challenges associated with the analysis of large and heterogeneous data sets, we developed the IntelliR data analysis platform. IntelliR provides standardized and automated data extraction, data summary, plotting, and statistical analyses for higher cognitive challenges, including the domains place learning, (multiple) reversal learning, extinction, episodic-like and working memory as well as executive functioning. The platform can be run upon individual challenges and can be modified with minimum R-programming knowledge for the analysis of additional modules. We also introduce conditional parameters, that are inaccessible in the default IntelliCage Analyzer software, to better assess cognition, exploration and repetitive behaviors. To validate the IntelliR platform and associated IntelliCage protocols, we confirmed that cognitive challenges are indeed learned by healthy mice, using 3 independent cohorts. Finally, we employed transgenic mice after sterile induction of pyramidal neuronal death in the *cornu ammonis* (19–21) and corroborated the ability to assess hippocampus-dependent cognitive function.

## METHODS

### Mice

All animal experiments were approved by the local animal care and use committee (LAVES, Niedersaechsisches Landesamt für Verbraucherschutz und Lebensmittelsicherheit, Oldenburg, Germany, license number 33.19-42502-04-18/2803) in accordance with the German animal protection law. Mice were housed in groups of up to 16 in a temperature (∼22°C) and humidity (∼50%) controlled environment, 12h light/dark cycle with food (standard food, Sniff Spezialdiäten, Germany) and water *ad libitum*. Cages were equipped with wood-chip bedding end nesting material (Sizzle Nest, Datesand, United Kingdom). Experimental mice were weaned at postnatal day 21 and separated by sex and genotype to avoid co-learning, inclusion effects, or aggressive behavior against possibly affected animals (1, 2, 22). Investigators performing animal experiments were unaware of group assignment (‘fully blinded’).

### Control cohorts

To avoid experiment-specific observations and to increase systematic variation to improve reproducibility (23), we analyzed data from 3 cohorts of female C57BL/6 mice aged 7-13 months with the following genotypes: heterozygous CNP-Cre (24) (n=16), heterozygous NexCreERT2 (19) (n=16), and wildtype (n=15).

### DTA cohort

Female C57BL/6 mice double heterozygous for Neurod6^tm2.1(cre/ERT2)Kan^ (‘NexCreERT2’, (19)) and Gt(ROSA)26Sor^tm1(DTA)Jpmb^ (‘Rosa26-eGFP-DTA’, (25)) (n=13), a tamoxifen inducible diphtheria toxin chain A allele, were used at the age of 7-8 months. Pyramidal cells were ablated as previously described (20) via intraperitoneal injections of 100mg tamoxifen (CAS#10540-29-1 T5648, Sigma-Aldrich) per kilogram body weight per day using a stock solution of 10mg/mL corn oil (C8267, Sigma-Aldrich) for 3 consecutive days. Control littermates received 10mL corn oil per kilogram body weight per day for 3 consecutive days. Place learning was started one week after the first tamoxifen injection. At this time point, pronounced pyramidal cell death and microgliosis were observed in the *cornu ammonis* region of male DTA littermates (n=4) after the same tamoxifen treatment. Loss of pyramidal neurons and hippocampal atrophy was confirmed histologically as previously described (20, 21) in tamoxifen treated mice (n=8), after behavioral phenotyping.

### Transponder placement

Mice were anesthetized with Avertin (2,2,2-tribromoethanol, Sigma-Aldrich, Taufkirchen, Germany, in ddH_2_O i.p. 300mg/kg body weight). After a small incision (2-3 mm) of the neck skin, one ISO standard transponder (12mm length, 2mm diameter, PM162-8; TSE Systems, Berlin, Germany) per mouse was implanted below the skin, enabling the IntelliCage system to individually identify each mouse. After closing the incision with suture and covering the eyes with protection salve, mice were put in their home cage, which was placed on a heating plate (∼38°C) to avoid hypothermia during the wake-up phase.

### IntelliCage Experiments

The structure and function of the IntelliCage paradigm has been previously described in detail (1, 2). In short, this setup allows observer-independent measurement of rodent behavior within a social environment. Being set inside a standard type IV rodent cage (20.5x55x38.5cm, Techniplast), the IntelliCage consists of 4 right-angled triangular conditioning chambers (“corners”, 15x15x21 cm, Fig.1A) and a food hopper with *ad libitum* access. The system provides 3 basic readouts for each conditioning chamber (Fig.1B): visits (ring radiofrequency identification antenna within the tube for recognizing individual mice via detection and readout of the implanted transponder), nose pokes (mouse disrupts light barriers in front of doors) and licks (number of contacts with lickometer). The IntelliCage system includes software to design experiments (when and where do animals get access to drinking nipples), online monitoring and collection of data. Using this software, we designed 3 challenges to test different cognitive abilities of mice. In between the challenges, mice received free access to water in all corners for 24h. During these extinction days, the health status and body weights of mice were assessed and IntelliCages were cleaned. Mice were excluded from the IntelliCage experiment, if they performed <100 licks/day on 2 consecutive days, if their health status deteriorated or if their body weight dropped.

**Figure 1:**
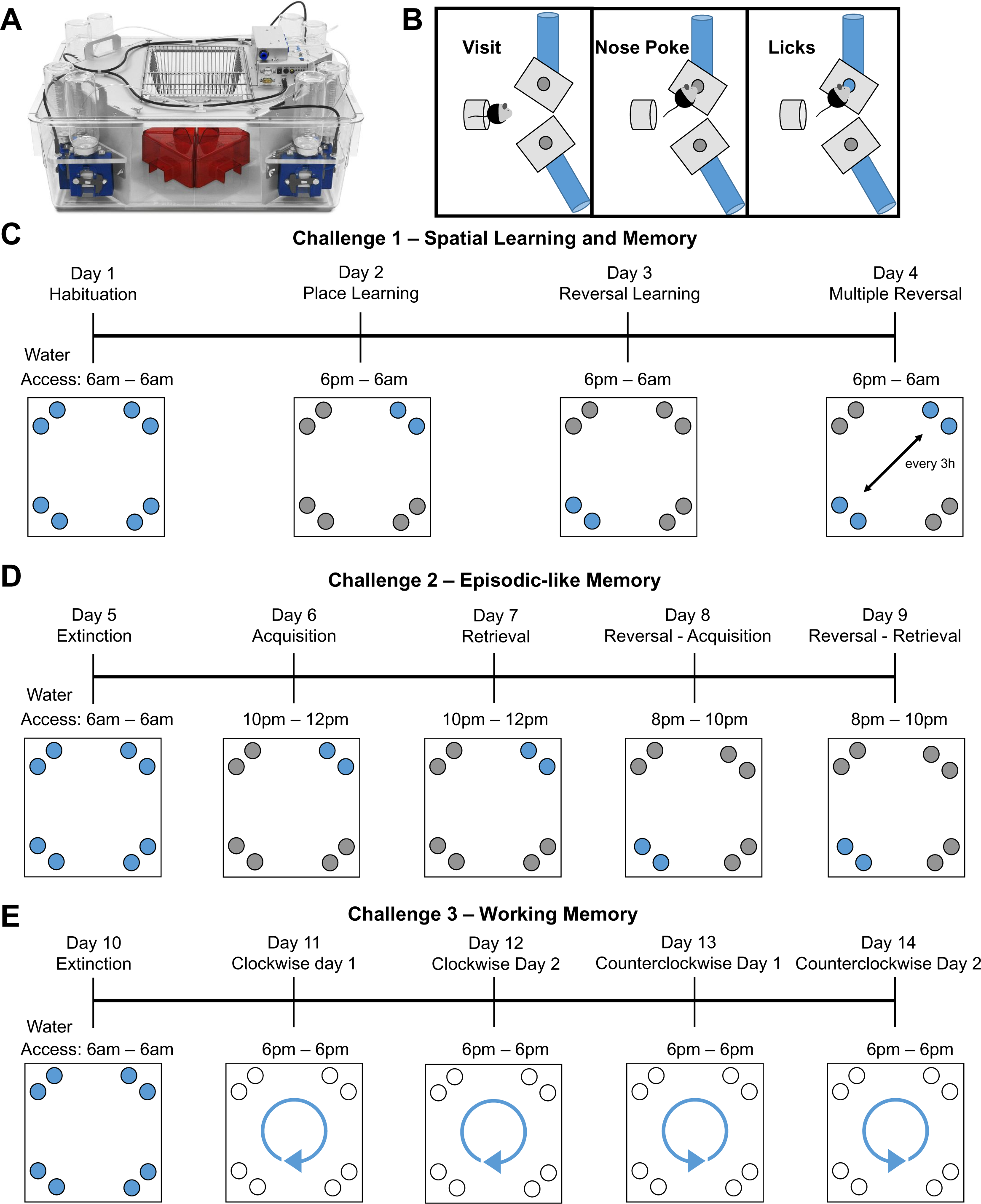
Descriptive overview of the IntelliCage setup and programmed challenges. **(A)** IntelliCage system for cognitive challenges of up to 16 mice. Conditioning corners allow access to water bottles and are equipped with sensors to detect presence and track behavior of mice within the corners. **(B)** Graphical description of the 3 basic behavior readouts provided by the system. Presence of mice within the tube leading to the water bottles (*visit*). Mice touching the door blocking the access to a water bottle with their noses (*nose poke*). After this, the door opens and the mice have access to water bottles and are able to drink (*licks*). **(C-E)** Graphical outline of the programmed challenges representative for one group. **(C)** On day 1 (d1), mice have free access for 24h to all water bottles in the corners (blue circles) to habituate to the new environment and task to get water. On d2, water access is only available in the dark phase (active phase) and one specific corner per group (*place learning*). On d3, water access only in the dark phase and the corner diagonally opposite of the previously correct corner (*reversal learning*). On d4, during dark phase, correct corner for water access switches every 3 hours between the 2 previously correct corners (*multiple reversal learning*). **(D)** After one day with free access to water for 24h (*extinction*, d5), challenge 2 starts with access to water from 10pm-12pm in the first of the previously learned corners (*episodic like memory – acquisition*). On d7, repetition of the settings from d6 (*episodic like memory – retrieval*). On d8, water access shifts to 8pm-10pm and to the corner diagonally opposite of the formerly correct corner (*episodic like memory reversal – acquisition*). On d9, repetition of the settings from d8 (*episodic like memory reversal – retrieval*). **(F)** After another day with free access to water for 24h (*extinction*, d10), challenge 3 tests for working memory via the patrolling paradigm. On d11 and d12, water access is granted during active phase and correct corners switch clockwise after each correct drinking attempt (nose poke). The first correct corner is defined by the first drinking attempt of the mice. On d13 and d14, correct corners switch counterclockwise after each correct drinking attempt. First correct corner is defined by the last correct drinking attempt during clockwise patrolling.

#### Challenge 1 – Spatial Learning and Memory

(Fig.1C): Mice were placed into the IntelliCage with free access to all water bottles when performing a nose poke (habituation) during the first light (6am-6pm) and dark phase (6pm-6am). Starting with the second light phase, water access was restricted to the active phase (dark phase, 12h) of the animals. Based on their transponder IDs, mice were divided into 4 groups per cage. During the second dark phase, each group was assigned to a single corner, where a nose poke would lead to water access. In the other 3 corners, doors would not open after performing a nose poke (*place learning*). In the third dark phase, each group was allocated the corner diagonally opposite of the previously correct corner (*reversal learning*). During the fourth dark phase, the correct corner for each group switched every 3 hours between the 2 formerly correct ones (*multiple reversal learning*).

#### Challenge 2 - Episodic-like Memory

(ELM, Fig.1D): This challenge is based on the episodic-like memory test developed by Dere and colleagues (1) and was modified to include a reversal of the where (corner) and when (time) component. The challenge starts with access to water in all corners after performing a nose poke. During the second dark phase, each group was allotted to the same corner that was rewarded during the place learning task. Water access was only granted between 10pm-12pm (ELM – acquisition). On the following dark phase, the same corner was rewarded with water access during the same time window to retrieve the previously learned task (ELM – retrieval). On the following 2 dark phases, the correct corner was again changed to the one diagonally opposite of the previously rewarded corner and the time window was shifted forward to 8-10pm (ELM reversal – acquisition and retrieval). The second time window for ELM reversal was specifically chosen to be set before the first one to avoid mice trying to get water access by extending their attempts into a new time window afterwards. Therefore, this task had to be learned anew and could not be fulfilled simply based on the first ELM task.

#### Challenge 3 – Working Memory

(Fig.1E): This challenge was programmed to change the correct corner after each correct nose poke in a clockwise pattern for 48h (*patrolling day 1 &* 2), followed by a switch to a counterclockwise pattern for the following 48h (*patrolling day 3 & 4*). The challenge starts with access to water in all corners after performing a nose poke during the first light and dark phase. To increase motivation and keep the start of cognitive testing consistent between challenges, mice were water deprived for 12h during the second light phase. After the beginning of the patrolling task in the second dark phase, all corners would give access to water if a nose poke is performed. The first nose poke of each individual mouse allocated this corner as the starting point for this challenge and therefore activated the patrolling protocol. After the experiment was stopped, the animals were transferred back to their home cages with food and water *ad libitum*.

### IntelliR Software

The present work introduces IntelliR, an easy-to-use platform for extracting relevant variables from the challenges described above via an R-script (26). It takes the form of a shiny-based (27) web interface, where users enter basic details from their experiment and receive processed variables, along with an initial statistical assessment and ggplot2 (28) figures that can be used for discussions. A full description of the required information and file organization, including tested package versions, is available in the provided documentation.

### Data Analysis

In addition to the basic measurements described in the IntelliCage Experiments section (i.e., visits, nose pokes, licks), IntelliR provides 9 main additional measurements calculated for the previously outlined challenges. However, the basic R script can be used and adapted to any IntelliCage design after minimal changes. The first pair of measurements evaluates the purpose of the visit executed by the mice. Exploratory visits are visits to corners where the mouse does not do a nose poke, in contrast to drinking attempts, which are visits where at least one nose poke was done. If more than one nose poke is performed in a single visit to a corner, it is possible that enhanced impulsivity with repetitive behavior has occurred. To quantify the frequency and intensity of potentially repetitive behaviors, the software extracts both the number of repetitive events, defined by the number of visits with multiple nose pokes (repetitive behavior frequency), as well as the number of repeated nose pokes (repetitive behavior total) as proxy of intensity. Of note, the repetitive behavior total can be triggered more than once in a single visit if, after a lick is registered, the mouse continues to make nose pokes without further licks. For example, 10 uninterrupted nose pokes in a single visit would result in a repetitive behavior frequency of 1 and a repetitive behavior total of 9.

The other 5 measurements are directly connected to the challenges and are named place error (PE), challenge error (CE), time error (TE), direction error (DE), and cognition index (CI). The first 4 reflect the specifics of the challenges, while CI compiles them to assess the performance of the mice. It is important to highlight that no single challenge contains all errors. The most basic error calculated is PE, which represents the ratio of drinking attempts made by a mouse in the incorrect corner in relation to all drinking attempts made. It is calculated for all challenges using the following formula:

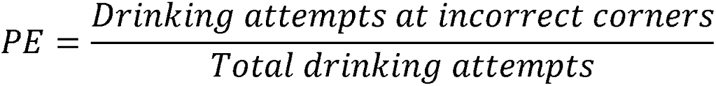

After the first day of each challenge, mice may remember the previously rewarded corner and be particularly drawn to it. CE is calculated in a similar fashion to PE, but takes into account whether the corner was rewarded in the previous challenge immediately before. It is calculated for Reversal Learning, Multiple Reversal Learning, Episodic-like Memory 2 Acquisition, and Episodic-like Memory 2 Retrieval. For challenges where CE is calculated, the formula for CE is:

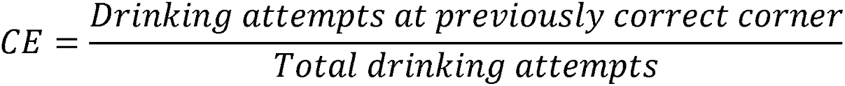

Challenge 2 integrates a time component that is also evaluated by IntelliR. TE is calculated for all 4 days and represents the proportion of drinking attempts made outside of the allotted time window. The formula for TE is:

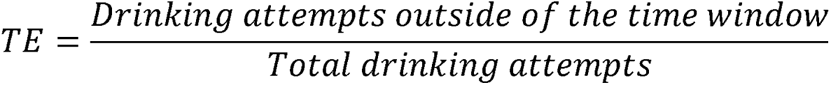

The last error, DE, is exclusive to days 3 and 4 of Challenge 3. On days 1 and 2, mice had to move clockwise to be able to continue drinking. On days 3 and 4, the movement changes to counterclockwise. DE evaluates how many times the animals go to the corner directly clockwise of the most recently rewarded corner. For example, if the rewarded corner was number 2 and the mouse makes drinking attempts in corner 3, it exhibits a DE. Drinking attempts in corners 2 or 4 do not contribute to it. The formula for DE is:

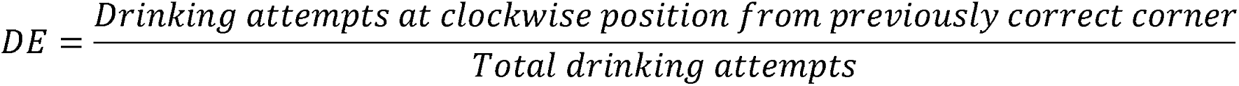

As each challenge can comprise different types of errors, the CI is calculated to summarize the cognitive performance of each mouse through the challenge. Errors are taken into account according to their severity, with the most basic one (i.e., PE) being ′punished′ the most. The specific calculation for each challenge is available in Table 1.

**Table 1:** Weights allotted to each type of error when calculating CI for individual challenge days.

| Challenge | Place Error (*adj.*) | Challenge Error | Time Error | Direction Error |
| --- | --- | --- | --- | --- |
| Place Learning | 100% | - | - |  |
| Reversal Learning | 75% | 25% | - |  |
| Multiple Reversal Learning | 75% | 25% | - |  |
| ELM Acquisition | 75% | - | 25% |  |
| ELM Retrieval | 75% | - | 25% |  |
| ELM Reversal Acquisition | 50% | 25% | 25% |  |
| ELM Reversal Retrieval | 50% | 25% | 25% |  |
| Working Memory Clockwise Day 1 | 100% | - | - |  |
| Working Memory Clockwise Day 2 | 100% | - | - |  |
| Working Memory Counterclockwise Day 1 | 75% |  | - | 25% |
| Working Memory Counterclockwise Day 2 | 75% | - | - | 25% |

As CE and DE are part of the overall PE, the place error rate had to be adjusted (PE_adj_) for the calculation of the cognition index using the following formula:

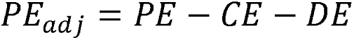

For the 4 types of errors, higher values indicate worse performance. However, for an index reflecting cognition, this seemed counter-intuitive and was therefore inverted. The overall CI formula is shown below:

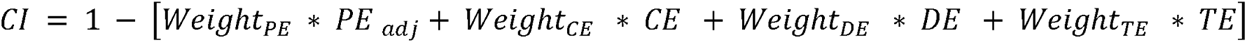

For evaluating our challenges, learning curves were calculated for all. For each task and mouse, an ordered sequence of drinking attempts was extracted and classified as PE or not. The sequence of drinking attempts and cumulative successful drinking attempts were then used to create learning curves by linear regression. A dummy group, representing by-chance-performance, was created using a slope of 0.25.

### Statistical Analysis

The current version of IntelliR provides statistical analysis for all calculated variables for both 2 or 4 groups. The statistical pipeline was made using R 4.3.0 (26) and base functions were used unless otherwise specified. Normality was tested through the Shapiro-Wilk test and equality of variance using Levene’s test from the car R-package (29).

When testing 2 groups, a variable with a parametric distribution is evaluated through a Two Sample t-test, with or without Welch’s correction. Effect sizes are calculated with Cohen’s d using the effectsize R-package (30). For non-parametric variables, Wilcoxon-Rank Sum Test is used, and a note is made to indicate whether or not the variable had equal variance. Effect sizes are calculated with Cliff’s delta using the effectsize R-package (30).

When 4 groups are specified, normality and equality of variance are calculated as described above. For variables with a parametric distribution, One-Way ANOVA is used taking into account the result of Levene’s test. Effect sizes are calculated using the Omega Squared test using the effectsize R-package (30). Post-hoc comparisons are done using the pairwise.t.test function (paired = FALSE) with Bonferroni correction for equal variances, while for unequal variances, the Games Howell Test from the rstatix R-package was used, which employs Tukey’s studentized range distribution for computing the p-values (31). In both cases, effect sizes are calculated with Cohen’s d using the effectsize R-package (30).

For variables with non-parametric distributions and equal variances, the Kruskal-Wallis Rank Sum Test is used. The epsilon-squared effect size is calculated with the following formula, where H is the Kruskal-Wallis test statistic and n is the sample size:

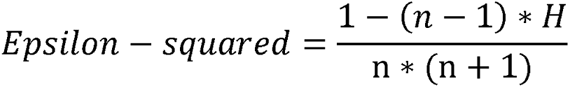

Post-hoc statistics are calculated using Dunn’s Test from the dunn.test R-package (32) and their effect sizes using Cliff’s delta from the effectsize R-package (30).

For non-parametric distributions with unequal variances, an Asymptotic K-Sample Fisher-Pitman Permutation Test is calculated using the coin R-package (33). Unfortunately, no adequate effect size test was found for this case. Post-hoc comparisons are performed using simultaneous tests for general linear hypothesis with multiple comparisons of means using the glht function of the multcomp R-package (34). Effect sizes are calculated using Hedges’ g through the emmeans R-package (35).

For the learning curves, global comparisons are calculated using analysis of covariance (ANCOVA), with specific group comparisons calculated through estimated marginal means contrasts using the emmeans R-package (35).

## RESULTS

### The IntelliR pipeline

To standardize and speed up the analysis of heterogeneous IntelliCage datasets, we developed the IntelliR pipeline along with 3 comprehensive cognitive challenges that assess spatial learning and memory, episodic-like memory, and working memory in a patrolling task (Fig.1). Furthermore, all challenges include reversal tasks to measure cognitive flexibility in assessed domains. IntelliR has an easy-to-use interface that within a few clicks and minutes extracts relevant information from IntelliCage experiments. The code is provided as an R-script and its basic structure can be easily modified to account for different challenges and experimental settings.

### Mice are able to learn all cognitive challenges provided by IntelliR

To confirm that mice perform as hypothesized and properly learn the cognitive tasks, we calculated learning curves of each task for 3 independent cohorts of healthy female mice tested in the IntelliCages (Fig.2). Comparing the learning curves with the average by-chance-performance using estimated marginal means contrasts revealed that all 3 independent cohorts successfully learned all eleven cognitive challenges as indicated by significant above-chance performance (all p<0.0001). Only a small number of individual mice failed to learn single tasks as indicated by similar to or worse than by-chance-performance (Fig.2).

**Figure 2:**
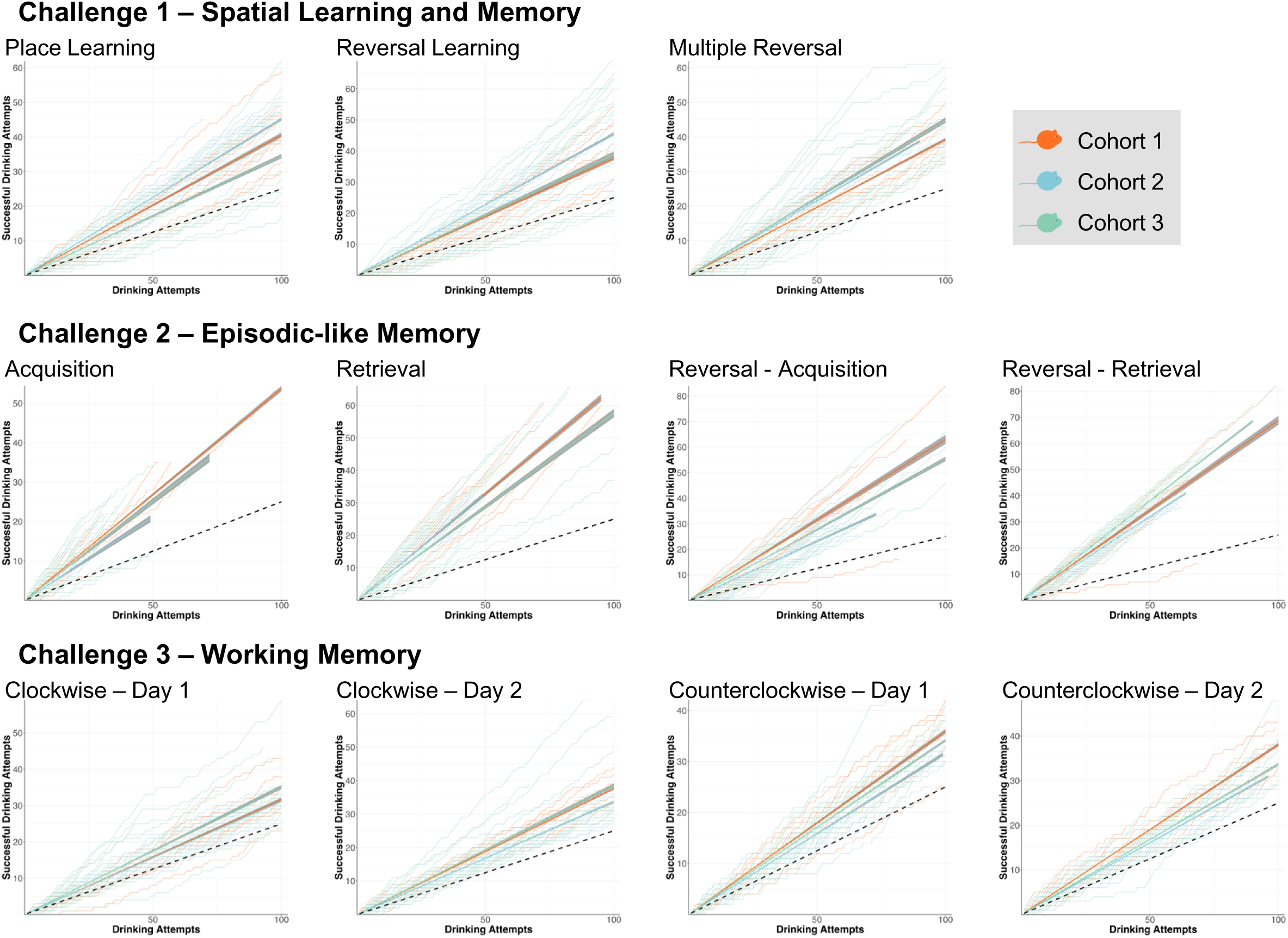
Learning performance of 3 independent cohorts of healthy female mice in individual cognitive tasks. Cumulative successes at each drinking attempt were plotted for individual mice (faint lines) and group performances calculated by linear regression (bold lines; 95% CI depicted as a shade) were compared to the average by-chance-performance (dashed line) through estimated marginal means contrasts. All 3 independent cohorts of mice successfully learned all cognitive tasks as indicated by significant above-chance performance (all p<0.0001). Only a small number of individual mice failed to learn some of the tests as indicated by close to or below chance performance. Global comparisons were calculated using analysis of covariance (ANCOVA), with specific group comparisons calculated through estimated marginal means contrasts.

### Assessing hippocampal dysfunction using IntelliR

To test whether the cognitive challenges are sensitive enough to detect hippocampal dysfunction, we utilized transgenic DTA mice in which pyramidal cell death can be acutely induced via tamoxifen (19–21). Pronounced neuronal cell death and microgliosis in the *cornu ammonis* region at the time of IntelliCage testing was confirmed using male DTA littermates, that were treated in parallel to the behavior cohort (Fig.3A-B). Loss of pyramidal neurons and hippocampal atrophy in DTA mice of the behavior cohort was confirmed histologically after behavioral phenotyping (Fig.3C).

**Figure 3:**
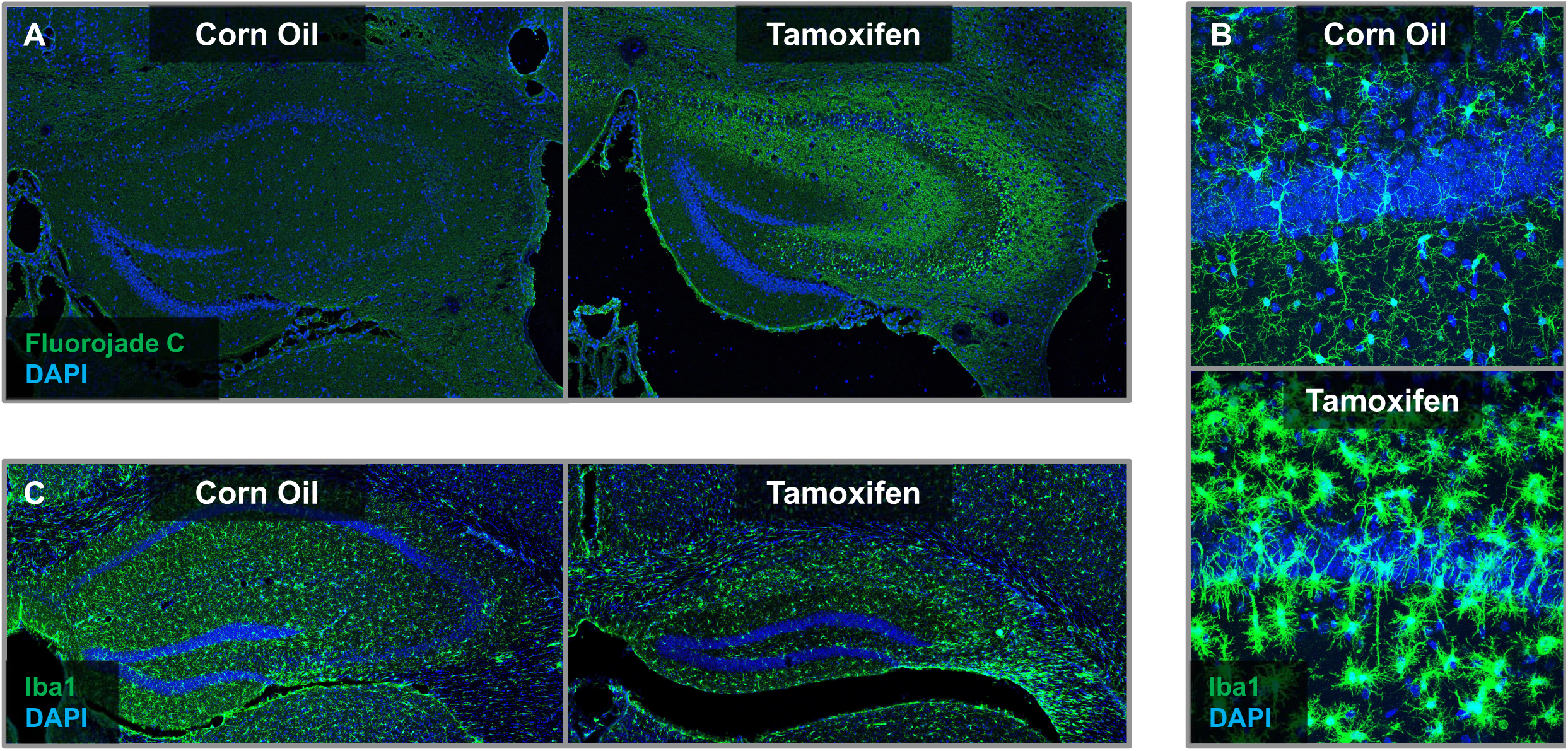
Histopathological consequences of tamoxifen induced diphtheria toxin A (DTA) expression in hippocampal pyramidal neurons. **(A)** Fluorojade C and DAPI staining of male DTA mice showed prominent neuronal degeneration in the cornu ammonis region 1 week after 3x tamoxifen injections as compared to DTA mice injected with 3x corn oil. **(B)** Iba1 and DAPI staining of hippocampal sections from mice presented in (A) show clear microgliosis upon tamoxifen induction. **(C)** Microgliosis and prominent hippocampal atrophy in tamoxifen treated DTA mice, that were used to validate the IntelliR pipeline. Eight mice per group (50%) were randomly selected and perfused for histological examination after the IntelliCage based phenotyping. For details on the immunohistochemical procedures see Wilke et al 2021 (20, 21).

The learning performance of DTA mice was evaluated after induction of hippocampal pyramidal cell death by tamoxifen, and compared to healthy corn oil treated littermate controls (Fig.4). While control mice showed a significant above-chance performance in all tests (p<0.0001), DTA mice failed to learn clockwise patrolling during the first day and showed significantly slower learning rates than controls (p<0.05) in all tests except place learning and ELM retrieval. Furthermore, we assessed the performance of mice with respect to individual cognitive components, namely the spatial component using the classical place error (Fig.5A), the contribution of sequential testing (extinction and cognitive flexibility) using the novel challenge error (Fig.5B) and direction error (Fig.5C), the temporal component via time error (Fig.5D) and summarized these diverse error types in a weighted cognition index (Fig.5E). In comparison to healthy corn oil controls, tamoxifen induced DTA mice showed a significantly higher rate of place errors in the first 24h of both clockwise and counterclockwise patrolling, whereas place and reversal learning, as well as place errors in the episodic-like memory test were not significantly altered (Fig.5A). Analysis of challenge errors revealed challenge-dependent differences in extinction and cognitive flexibility. While tamoxifen induced mice made significantly less challenge errors in the multiple reversal learning trial, they showed significantly higher challenge errors in the ELM reversal acquisition day (Fig.5B). Extinction of the clockwise patrolling task was similar between tamoxifen and corn oil treated DTA mice and completed within the first day of counterclockwise patrolling (Fig.5C). Similarly, learning of the temporal component of the episodic-like memory test was comparable between tamoxifen and corn oil treated mice (Fig.5D). Taking the different types of errors into account, the cognition index shows, that within each challenge, mice improved their cognitive performance with time (Fig.5E). Furthermore, the cognition index reveals worse cognitive performance in the patrolling task indicating impaired working memory in mice upon ablation of hippocampal pyramidal neurons (Fig.5E).

**Figure 4:**
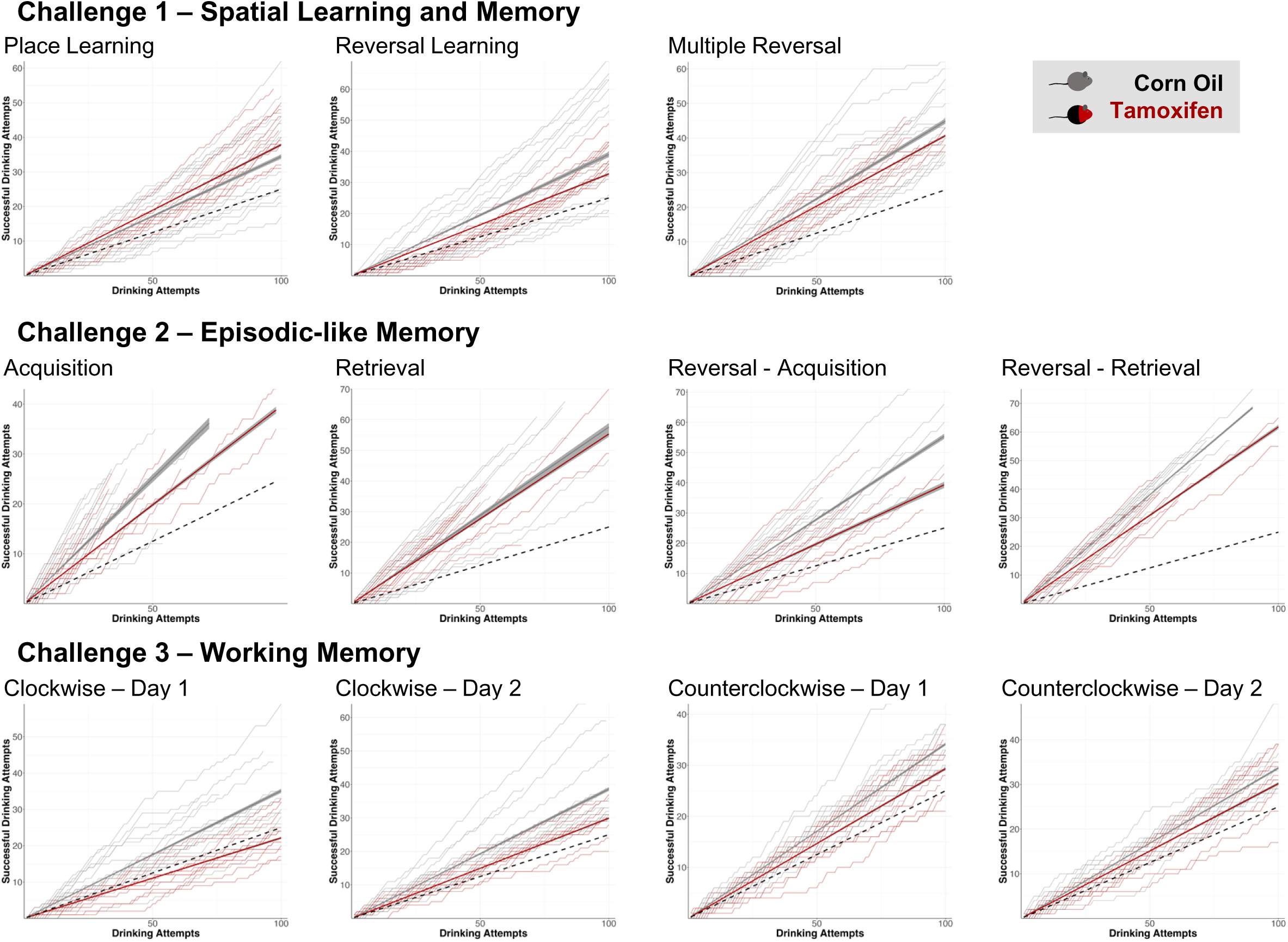
Learning performance of the DTA cohort. To test if the cognitive challenges are sensitive enough to detect hippocampal dysfunction, the learning performance of DTA mice was tested after induction of hippocampal pyramidal cell death by tamoxifen and compared to corn oil treated healthy littermate controls. Cumulative successes at each drinking attempt were plotted for individual mice (faint lines) and group performances, calculated by linear regression (bold lines; 95% CI depicted as gray shadows), were compared to the average by chance performance (dashed line) and between groups through estimated marginal means contrasts. While control mice showed a significant above-chance performance in all tests (p<0.0001), DTA mice failed to learn clockwise patrolling during day 1 and showed significantly slower learning rates (p<0.05) than controls in all tests except place learning and episodic-like memory retrieval. Global comparisons were calculated using analysis of covariance (ANCOVA), with specific group comparisons calculated through estimated marginal means contrasts.

**Figure 5:**
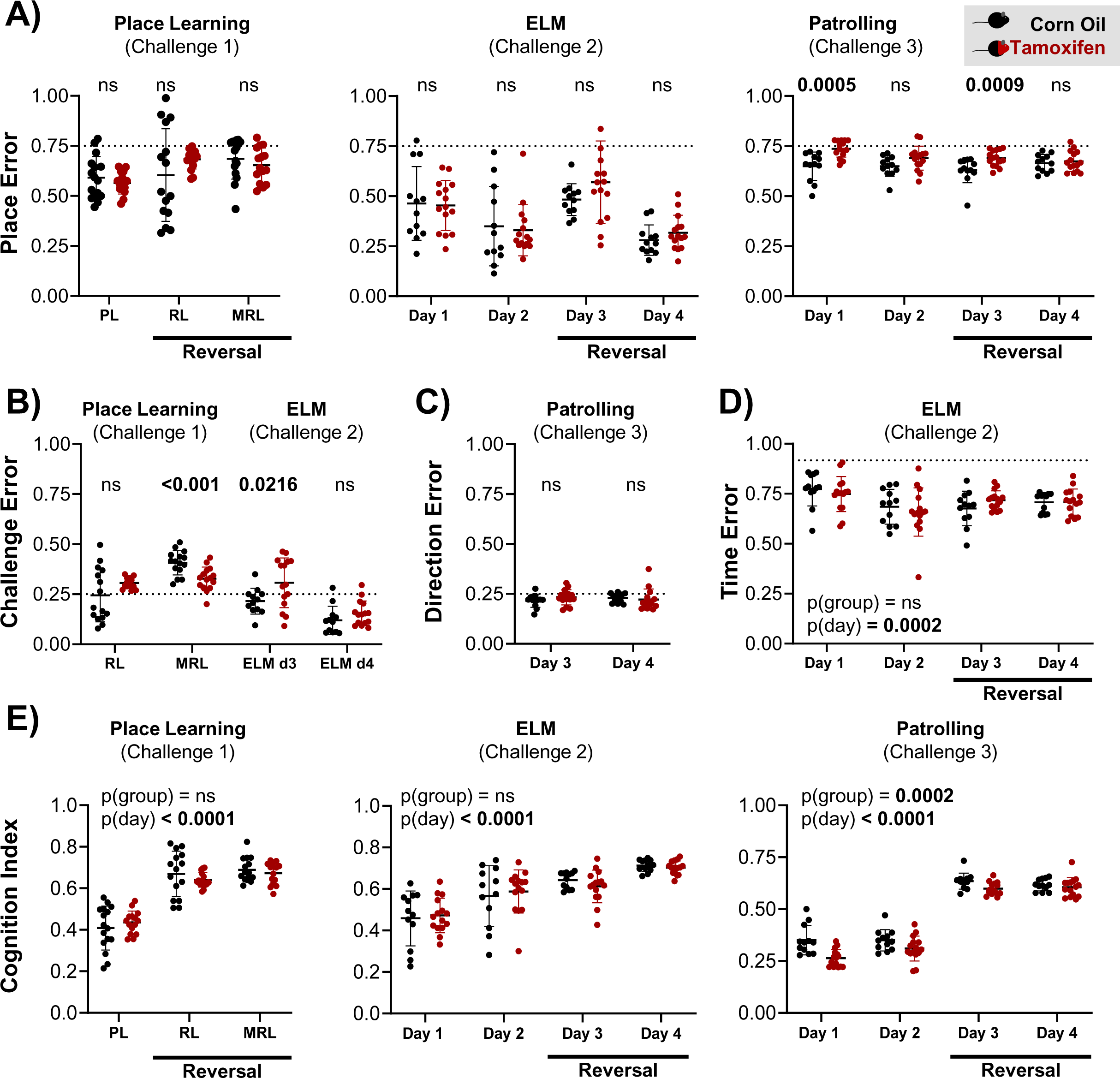
Cognitive performance of tamoxifen induced (red) versus corn oil treated (black) DTA mice. **(A)** Place errors are plotted to evaluate the spatial component of all cognitive tests. DTA mice show significant deficits during the first day of clockwise and counterclockwise patrolling respectively. **(B, C)** Challenge errors **(B)** and direction errors **(C)** are presented for all challenges involving a reversal task to evaluate extinction of the previous task and cognitive flexibility. The dotted lines indicate the average by-chance-performance (3/4 corners are incorrect). **(D)** Time errors are plotted to assess learning of the temporal component of the episodic-like memory test. Average by-chance-performance is indicated by the dotted line (22/24h are incorrect). **(E)** To assess the overall cognitive performance of mice across all challenges, the cognition indices are plotted for all challenges. Taking the different types of errors into account, the cognition index shows that within each challenge mice significantly improved their cognitive performance with time (all p<0.0001, repeated measures ANOVA for Challenge 1 and Friedman’s two-way test for Challenges 2 and 3). Furthermore, the cognition index shows a significantly worse cognitive performance in the patrolling task in mice upon hippocampal pyramidal cell ablation (p=0.0002, Friedman’s two-way test). Data presented as mean ± standard deviation. Pairwise comparisons in **A-D** calculated as described in the method section.

### Assessing basic mouse behaviors using IntelliR

In addition to the automated cognitive phenotyping, IntelliR provides users with parameters to monitor more basic mouse behavior, such as overall (circadian) activity, exploratory behavior, and repetitive behaviors. These parameters are invaluable for the assessment of the overall health and well-being of mice as they are continuously recorded in a social home cage environment (2). Analyzing the number of visits of 3 independent cohorts of healthy female mice during Challenge 1 in our IntelliCage paradigm demonstrated that the majority (85.15%) of visits occurred during the dark as compared to the light phase (580.5±156.8 versus 128.3±59.3, p<0.001), highlighting the usefulness of visits as proxy for general activity.

Furthermore, an analysis of the number of exploratory visits per test day showed a prominent decrease during the same period (Friedman Rank Sum test p<0.001), indicating that exploratory visits are indeed a good proxy of exploratory behavior. While we did not observe differences in general activity, exploratory behavior, or number of repetitive events between tamoxifen and corn oil treated DTA mice throughout the challenges (Fig. 6A-C), these parameters can be useful for mouse models with more pronounced motor or stereotypic phenotypes, such as models for Huntington disease, Parkinson disease, multiple sclerosis, or autism spectrum disorder (36–39).

**Figure 6:**
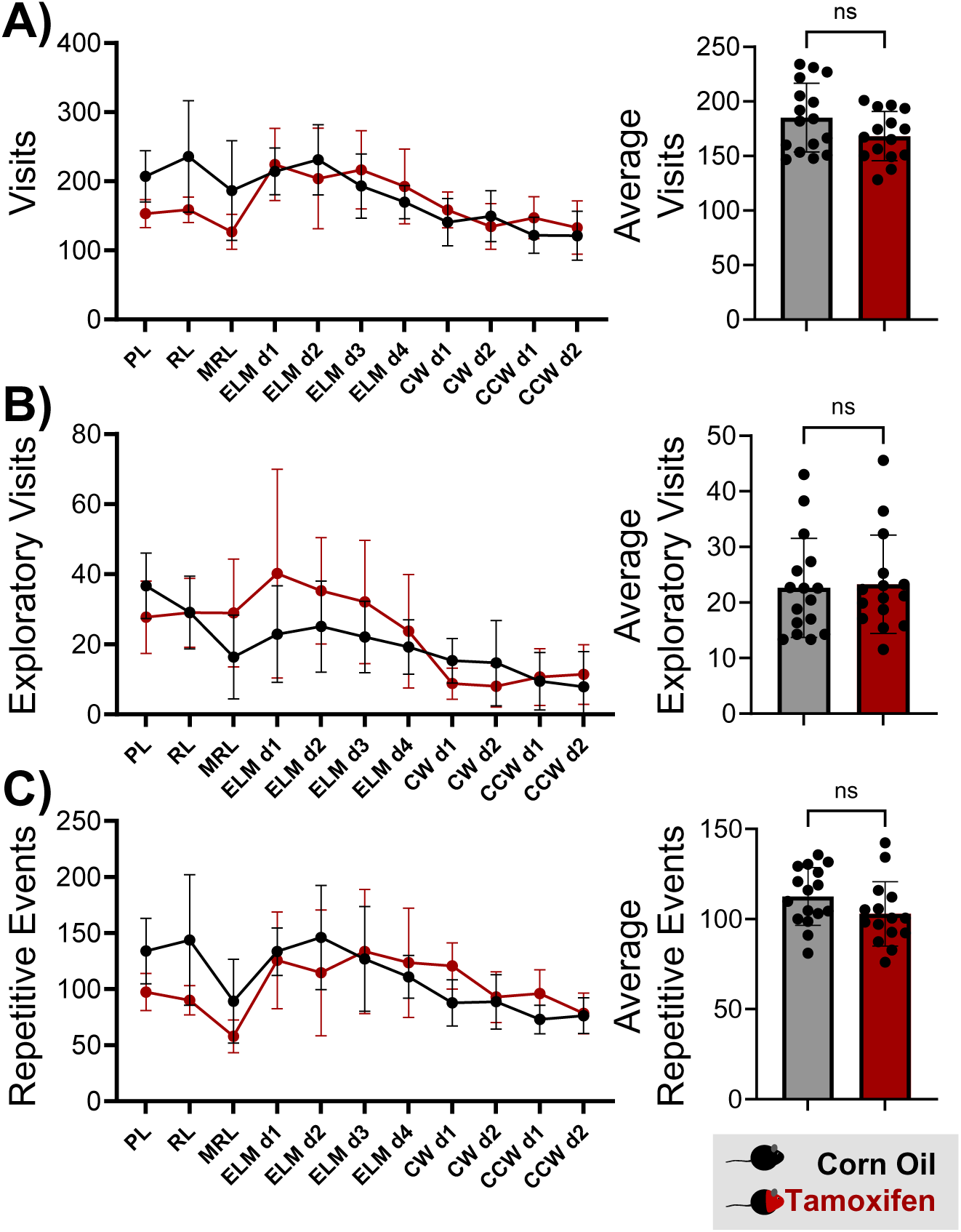
Monitoring basic activity related mouse behavior using IntelliR. Total number of corner visits **(A)**, exploratory visits **(B)** and repetitive events **(C)** per test day (left graphs) and averages over all test days (right graphs) were used to evaluate general activity **(A)**, exploratory behavior **(B)** and repetitive behavior **(C)**. No differences were observed between tamoxifen (red) and corn oil (black) treated DTA mice. Data presented as mean ± standard deviation. Welch’s corrected two-sided t-tests were used to compare the averaged parameters.

## DISCUSSION

### Summary

IntelliR has been designed to offer a comprehensive pipeline for analyzing datasets generated by IntelliCages. As a tool, IntelliCages play a pivotal role in behavioral analysis of mice by enabling independent longitudinal testing in the social context of their home cage. IntelliR standardizes the analysis process, thereby allowing researchers to concentrate more on interpreting their results rather than the intricacies of data analysis.

Furthermore, IntelliR significantly simplifies the process of evaluating basic activity of mice, a crucial metric in assessing their well-being. The primary advantage of the IntelliCage is its ability to monitor and assess mouse behavior with a minimum of experimenter interference, thereby providing for the animals a more familiar and likely less stressful environment (2). Another advantage of the IntelliR is that it enables automated cognitive profiling of mice as they are exposed to a diverse set of cognitive challenges and offers a number of additional parameters, which are inaccessible in the IntelliCage Analyzer software.

### Validity of IntelliCage Challenges

The selected challenges were based on existing IntelliCage test designs that assess simple place learning and spatial memory (1), episodic-like memory (1), as well as reference memory and spatial working memory in a clockwise patrolling task (10, 40, 41). For all challenges, reversal tasks were included to evaluate cognitive flexibility in the assessed domains. In addition, we included days without any cognitive task and with unrestricted access to water, that allow the experimenter to clean the cages, assess animal welfare, and appraise extinction of learned behavior.

To independently validate that healthy mice are able to learn these challenges within the allocated testing times, we tested 3 independent cohorts of female mice and introduced systematic variation to enhance reproducibility and avoid experiment-dependent observations (23). We demonstrated that our proposed challenges can be successfully learned by healthy mice, with their performance significantly exceeding random chance. Furthermore, we evaluated the sensitivity of these cognitive tasks for measuring hippocampal dysfunction by testing DTA mice after ablation of hippocampal pyramidal neurons. Consistent with previous findings in this model (21) and mice with hippocampal lesions (9, 10), we did not observe differences in the IntelliCage-based place learning and reversal learning tasks. In contrast, the patrolling task revealed notable differences in our DTA model, which is consistent with previous findings in other mouse models of hippocampal dysfunction (10, 40), stressing the importance of this test for the assessment of disturbed behavioral patterns and neuropsychiatric symptoms.

### Measuring success in different cognitive tasks

A reliable and frequently used measurement of success in tasks with a spatial component in the IntelliCage is the **place error**. The place error indicates how often mice fail to visit or nose poke the correct corner, or place preference, which indicates how frequent mice visit or nose poke the rewarded corner (2, 12–15, 41–51). In contrast to previous studies that calculated place errors based on the number of corner visits or number of nose pokes, IntelliR provides error rates based on the number of drinking attempts. We defined drinking attempts as visits in which mice attempt to access water using the learned operant response (nose poke), thereby minimizing the influence of exploratory corner visits. To evaluate how well mice learn a particular time association in tasks with a temporal component, IntelliR provides a **time error**. Furthermore, sequential testing of mice in different cognitive tasks requires mice not only to learn the new rule but also not to follow the previous rules (52, 53). To appraise the ability of mice for behavioral extinction, IntelliR provides **challenge error and direction error**, that indicate how often a mouse follows the rules of the preceding task. To assess the global cognitive performance in all of these components, we introduced the cognition index. Comparing the cognition index of mice across different cognitive tasks revealed that healthy mice improved their cognitive performance with successive tests and outperformed DTA mice in the patrolling task.

### Validity of activity-related parameters

As the movement of mice is not directly tracked, the **general activity** in the IntelliCage is typically measured using the number of corner visits as proxy (2). Using this approach, a variety of researchers have replicated activity-related phenotypes observed in classical behavioral tests and different automated tracking systems in the IntelliCages (5, 37–39, 45, 54, 55). Furthermore, we recently confirmed that the number of corner visits in the IntelliCage correlates with the locomotor activity in an automated vibration-based phenotyping system (21). To show that corner visits are indeed a good proxy for activity, we analyzed the number of corner visits of healthy mice during the dark phase as compared to the light phase and found that approximately 85% of the corner visits occur in the active (dark) phase of mice, thereby confirming corner visits as proxy for activity and demonstrating the capacity to monitor circadian rhythms.

To assess **exploratory behavior** in the IntelliCage, we hypothesized, that visits to the conditioning corners, in which mice do not attempt to access water with a nose poke, are due to exploratory behavior and defined these visits as **exploratory visits**. This hypothesis is based on the assumption that mice quickly learn, that access to water is limited to the conditioning corners and requires an operant response, i.e., nose poke. Indeed, learning of the operant response and adaptation to the IntelliCage environment is typically achieved within one day, either during an initial habituation or free adaptation phase (11), or directly during a place learning task (1). As exploratory behavior decreases with increased familiarity to a novel environment such as the IntelliCage (11, 13, 41) we assessed the number of exploratory visits of mice during the exposure to the IntelliCage environment and found a significant decrease with time, indicating that exploratory visits can be used as proxy for exploratory behavior.

To assess the intensity and frequency of **repetitive behavior**, IntelliR provides the number of repetitive nose pokes and the number of repetitive events, both defined as uninterrupted bouts of at least 2 nose pokes. Although a similar approach has been described previously (56), the threshold for repetitive nose pokes and repetitive events may need to be adjusted. Careful validation with a mouse model, characterized by consistent repetitive, impulsive, or stereotypic behavior, will help achieve optimal sensitivity and specificity of this readout.

### Limitations

This study has limitations that can be broadly categorized into general and technical aspects. The research was conducted using multiple cohorts of healthy mice and one specific model. However, all the subjects were female and of a certain age.

Therefore, additional testing is necessary to determine whether the findings are applicable to male mice, mice with different phenotypes, and older mice. Specifically for the last two cases, severe movement limitations, such as hind limb paralysis in mice with experimental autoimmune encephalomyelitis, can interfere with their ability to successfully enter the conditioning corners. Furthermore, mice that fail to visit or drink in the correct corner, i.e., due to lack of motivation, cognitive impairments or anxiety, have to be removed from the IntelliCage due to water deprivation. In these cases, the experimenter may consider changing the reward to sucrose solution and providing *ad libitum* access to water outside of the conditioning corners (2).

A theoretical consideration to keep in mind when using the cognition index is that it assumes that all challenges are learned and that extinction of previously learned tasks requires trial and error, reflected by lower weight of challenge and direction errors as compared to place error. Testing 3 independent cohorts of healthy female mice confirmed this assumption.

The main technical limitation is that the current state of IntelliR requires that all challenges are performed and executed as described by us. These might not be ideal for different experimental settings and questions. We try to mitigate this by providing the design files for the challenges which can be modified. IntelliR is also available as an open R-script and can be changed for different settings with a moderate amount of knowledge.

In conclusion, we present an analysis pipeline in the form of an R-script, IntelliR, along with a set of challenges designed to provide a sensitive measure for higher cognition profiling. Despite its limitations, our work offers a standardized approach to behavioral valuations, upgrading the usability of IntelliCages, and likely contributing towards reproducibility in the field.

## DATA AVAILABILITY

The IntelliCage data that support the findings of this study are available from the authors upon request. Current IntelliR code, IntelliCage designer files, and manual, as well as future updates, are available from GitHub under https://github.com/vgastaldi/IntelliR.

## AUTHOR CONTRIBUTIONS

Supervision: HE, KAN

Funding acquisition: HE, KWM, KAN

Concept and design: VDG, MH, JBHW, SA, HE

Data acquisition/generation: VDG, MH, AR, JBHW, AFW, AN

Data analyses/interpretation: VDG, MH, JBHW, AR, SK, AFW, AN, HE

Drafting the manuscript: VDG, MH, JBHW, HE

Drafting display items: VDG, MH, JBHW, HE

Continuous critical input, review & editing: MJP, LY, YC, JACS, UJB, KWM, KAN, HE

**All authors read and approved the final version of the manuscript.**

## ACKNOWLEDGEMENTS

This work has been funded by the European Research Council (ERC) Advanced Grant to HE under the European Union’s Horizon Europe research and innovation programme (acronym *BREPOCI;* grant agreement No 101054369), as well as an ERC Consolidator Grant to KWM (acronym *ALTIBRAIN*; grant agreement No 101043416), in collaboraton with HE. Furthermore, the study has been fostered by the Max Planck Society and the Max Planck Förderstiftung. Research in the labs of HE and KAN is funded by DFG TRR-274/1 2020-408885537. KAN is supported by the Adelson Medical Research Foundation. VDG received support from the IMPRS-Genome Science PhD program.

## List of Abbreviations

CE: challenge error
CI: cognition index
ELM: episodic-like memory
DE: direction error
PE: place error
TE: time error.

